# Comparison of different linear-combination modelling algorithms for short-TE proton spectra

**DOI:** 10.1101/2020.06.05.136796

**Authors:** Helge J. Zöllner, Michal Považan, Steve C. N. Hui, Sofie Tapper, Richard A. E. Edden, Georg Oeltzschner

## Abstract

Short-TE proton MRS is used to study metabolism in the human brain. Common analysis methods model the data as linear combination of metabolite basis spectra. This large-scale multi-site study compares the levels of the four major metabolite complexes in short-TE spectra estimated by three linear-combination modelling (LCM) algorithms.

277 medial parietal lobe short-TE PRESS spectra (TE = 35 ms) from a recent 3T multi-site study were pre-processed with the Osprey software. The resulting spectra were modelled with Osprey, Tarquin and LCModel, using the same three vendor-specific basis sets (GE, Philips, and Siemens) for each algorithm. Levels of total N-acetylaspartate (tNAA), total choline (tCho), myoinositol (mI), and glutamate+glutamine (Glx) were quantified with respect to total creatine (tCr).

Group means and CVs of metabolite estimates agreed well for tNAA and tCho across vendors and algorithms, but substantially less so for Glx and mI, with mI systematically estimated lower by Tarquin. The cohort mean coefficient of determination for all pairs of LCM algorithms across all datasets and metabolites was 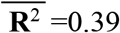, indicating generally only moderate agreement of individual metabolite estimates between algorithms. There was a significant correlation between local baseline amplitude and metabolite estimates (cohort mean 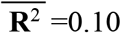).

While mean estimates of major metabolite complexes broadly agree between linear-combination modelling algorithms at group level, correlations between algorithms are only weak-to-moderate, despite standardized pre-processing, a large sample of young, healthy and cooperative subjects, and high spectral quality. These findings raise concerns about the comparability of MRS studies, which typically use one LCM software and much smaller sample sizes.

**Graphical Abstract:** 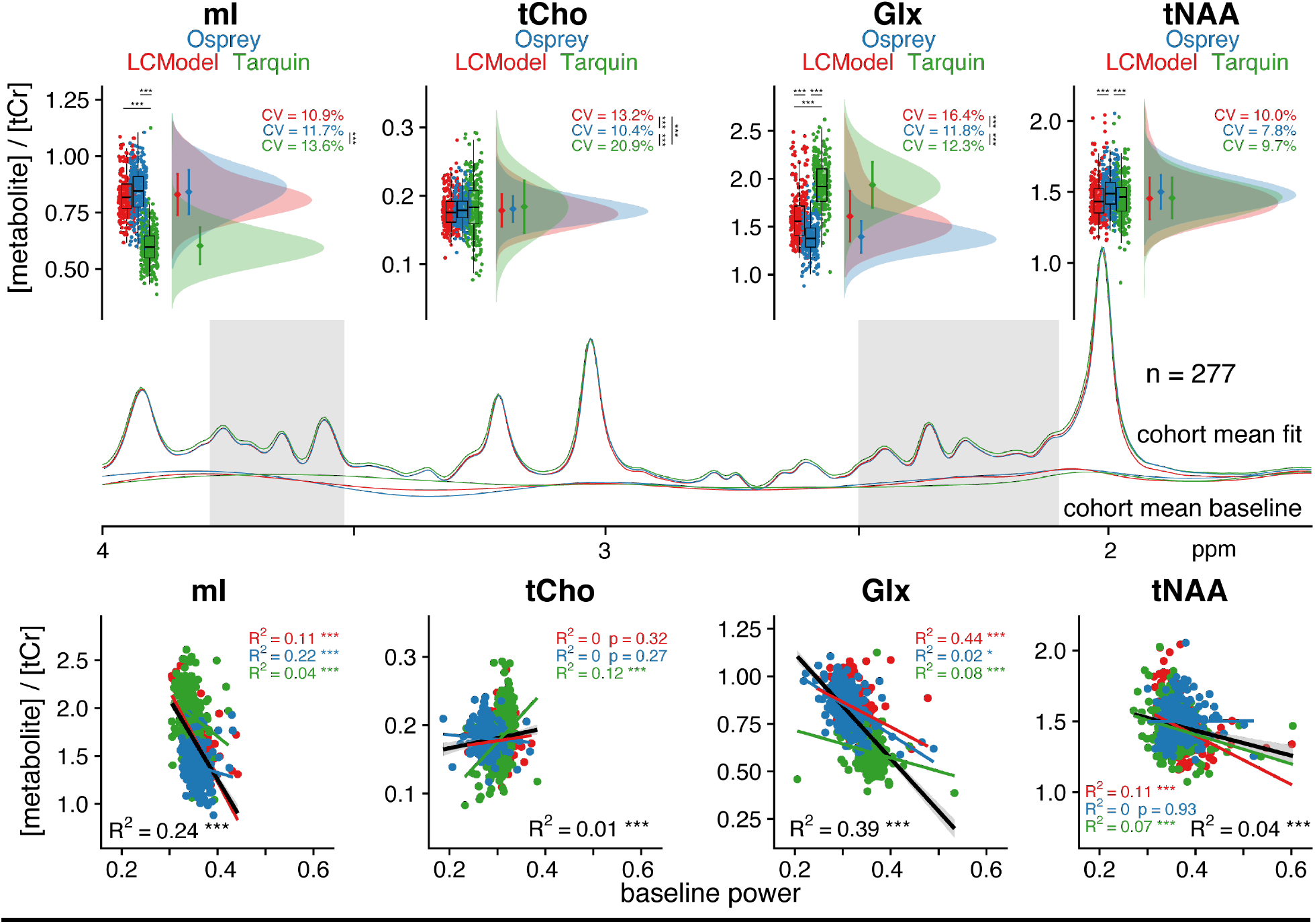

Three linear-combination algorithms (Osprey, Tarquin and LCMode) were used to quantify the levels of tNAA, tCho, mI, and Glx in 277 short-TE PRESS. Group means and CVs of metabolite estimates agreed well for tNAA and tCho, but substantially less so for Glx and mI with a cohort mean correlation coefficient of 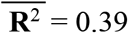, indicating moderate agreement between algorithms. These findings raise concerns about the comparability of MRS studies, which typically use one LCM software and much smaller sample sizes.

## Introduction

Proton MRS allows in-vivo research studies of metabolism^1,2^. Single-voxel MR spectra from the human brain are frequently acquired using PRESS localization^3^, and can be modelled to estimate metabolite levels. Accurate modelling is hampered by poor spectral resolution at clinical field strengths, and for short-echo-time spectra, metabolite signals overlap with a broad background consisting of fast-decaying macromolecule and lipid signals. Linear-combination modelling (LCM) of the spectra maximizes the use of prior knowledge to constrain the model solution, and is recommended by recent consensus^4^. LCM algorithms model spectra as a linear combination of (metabolite and macromolecular (MM)) basis functions, and typically also include terms to account for smooth baseline fluctuations.

Several LCM algorithms are available to quantify MR spectra (**Table 1** describes some of the most widely used: Osprey^5^, INSPECTOR^6^, Tarquin^7^, AQSES^8^, Vespa^9^, QUEST^10^, LCModel^11^). The implementations (open-source vs. compiled ‘black-box’), modelling approaches (modelling domain and baseline model), and their licensure practices are diverse. The most widely used algorithm is the LCModel implementation (accounting for ∼90% of the citations in Table 1), often considered as the gold standard, apart from it being the prototype LCM algorithm for MRS quantification with a pre-compiled ‘black-box’ implementation and a substantial price tag. Surprisingly few studies have compared the performance of different LCM algorithms. Cross-validation of quantitative results has almost exclusively been performed in the context of bench-marking new algorithms against existing solutions. In-vivo comparisons are often limited to small sample sizes, whether analyzing spectra from animal models^7,12,13^ or human subjects^7,8,12^. To the best of our knowledge, two exceptions compared the LCM performance of different algorithms in rat brain^14^ and human body^15^, respectively. Most studies report good agreement between results from different algorithms, inferring this from group-mean comparisons, or observing that differences between clinical groups are consistent regardless of the algorithm applied^14,16^. Correlations of estimates from different algorithms are rarely reported; however, a high correlation between LCModel and Tarquin results was found in the rat brain at ultra-high field^14^. Despite the fact that LCM has been used to analyze thousands of studies (**Table 1**), a comprehensive assessment of the agreement between the algorithms is lacking, and the relationship between the choice of model parameters and quantitative outcomes is poorly understood. To begin to address this gap, we conducted a large-scale comparison of short-TE in-vivo MRS data using three LCM algorithms with standardized pre-processing. While recent expert consensus recommends using measured MM background spectra, data for different sequences are not broadly available or integrated in LCM software. This manuscript investigates current common practice, and therefore all models included simulated MM basis functions as defined in LCModel. We compared group-mean quantification results of four major metabolite complexes from each LCM algorithm, performed between-algorithm correlation analyses, and investigated local baseline power and creatine modelling as potential sources of differences between the algorithms.

**Table 1.** Overview of linear-combination modelling algorithms. The domain (time TD or frequency FD) of modelling and the baseline model approach are specified. *Citations reported from Google Scholar on October 26, 2020.

| **Name** | **Modelling Domain,**<br>**Baseline approach** | **Cost** | **Code**<br>**Availability** | **Published** | **Cita-**<br>**tions\*** |
| --- | --- | --- | --- | --- | --- |
| Osprey | FD, spline baseline | free | open | 2020 | 1 |
| INSPECTOR | FD, 1<sup>st</sup>-order polynomial | free | open | 2018 | 0 |
| Tarquin | TD, smooth baseline | free | open | 2011 | 264 |
| AQSES<br>(jMRUI) | TD, spline baseline | free | closed | 2007 | 143 |
| Vespa | FD, wavelet baseline | free | open | 2006 | 75 |
| QUEST<br>(jMRUI) | TD, spline baseline | free | closed | 2004 | 35 |
| LCModel | FD, spline baseline | \$13,300 | closed | 1992 | 3518 |

## Methods

### Participants & acquisition

277 single-voxel short-TE PRESS brain datasets from healthy volunteers acquired in a recent 3T multisite-study^17^ were included in this analysis. Data were acquired at 25 sites (with up to 12 subjects per site) on scanners from three different vendors (GE: 8 sites with n = 91; Philips: 10 sites with n = 112; and Siemens: 7 sites with n = 74) with the following parameters: TR/TE = 2000/35 ms; 64 averages; 2, 4 or 5 kHz spectral bandwidth; 2048-4096 data points; acquisition time = 2.13 min; 3×3×3 cm^3^ voxel in the medial parietal lobe (**Figure 1A)**. The water suppression pulse bandwidth was 140 Hz for Philips, 50 Hz for Siemens, and 150 Hz for GE. Reference spectra were acquired with similar parameters, but without water suppression and 8-16 averages. No more acquisition parameters were specified (for more details, please refer to ^17^). Data were saved in vendor-native formats (GE P-files, Philips .sdat, and Siemens .dat). In the initial study^18^, written informed consent was obtained from each participant and the study was approved by local institutional review boards. Anonymized data were shared securely and analyzed at Johns Hopkins University with local IRB approval. Due to site-based data privacy guidelines, only a subset of these data (GE: 7 sites with n = 79; Philips: 9 sites with n = 100; and Siemens: 4 sites with n = 48) is publicly available^19^.

**Figure 1.**
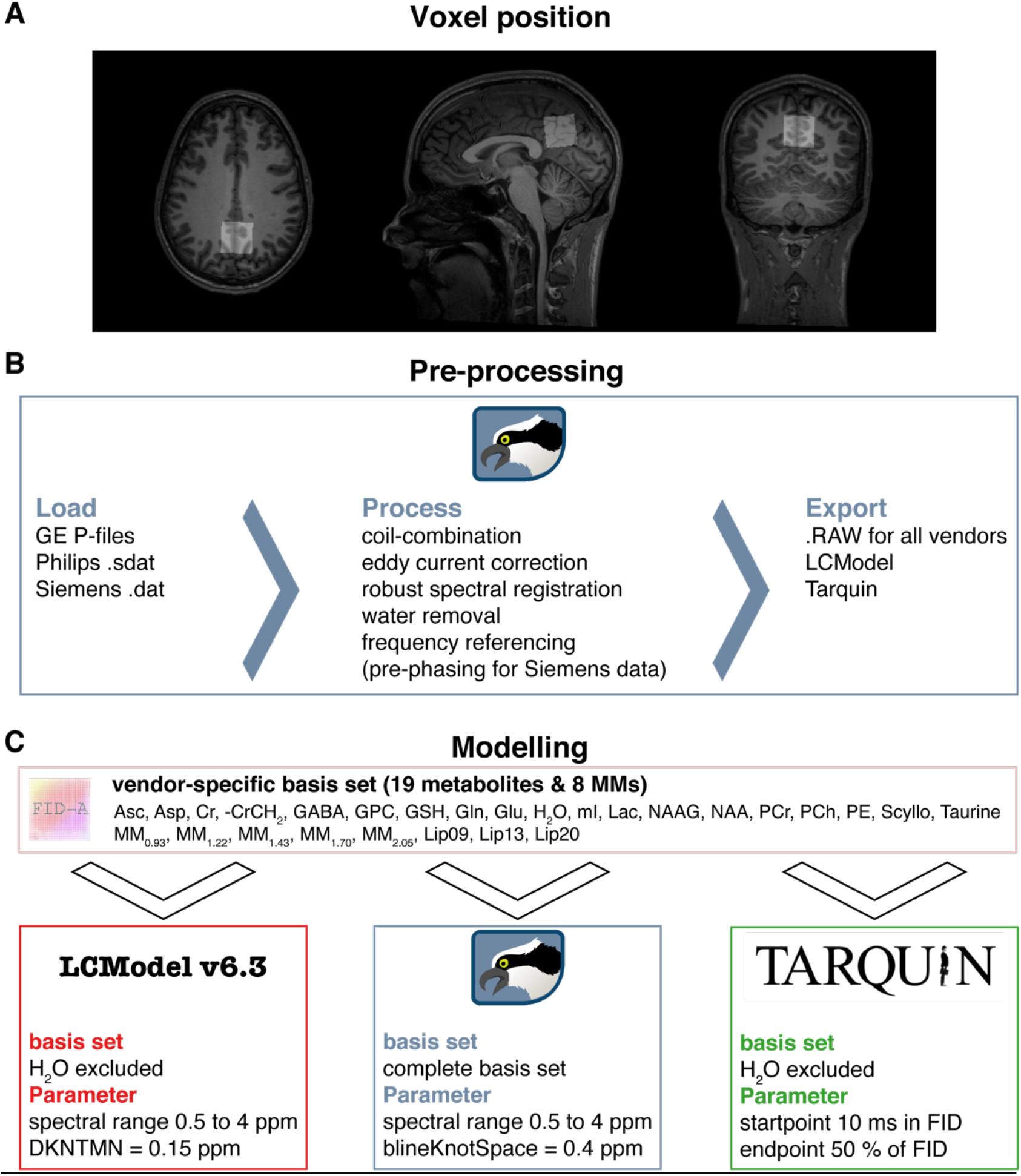
Voxel position and overview of the MRS analysis pipeline. (A) Representative voxel position in the medial parietal lobe extracted with ‘OspreyCoreg’ (B) Pre-processing pipeline implemented in Osprey including ‘OspreyLoad’ to load the vendor-native spectra, ‘OspreyProcess’ to process the raw data and to export the averaged spectra. (C) Modelling of the averaged spectra with details of the basis set and parameters of each LCM (LCModel, Osprey, and Tarquin).

### Data pre-processing

MRS data were pre-processed in Osprey^5^, an open-source MATLAB toolbox, following recent peer-reviewed pre-processing recommendations^2^, as summarized in **Figure 1B**. First, the vendor-native raw data were loaded, including the metabolite (water-suppressed) data and unsuppressed water reference data. Second the raw data were pre-processed into averaged spectra. Receiver-coil combination^20^ and eddy-current correction^21^ of the metabolite data were performed using the water reference data. Individual transients in Siemens and GE data were frequency-and-phase aligned using robust spectral registration^22^. The Philips data had been coil-combined by weighted combination with the complex coefficients obtained during the survey scan and averaged on the scanner without frequency-and-phase correction of the individual transients. After averaging the individual transients, the residual water signal was removed with a Hankel singular value decomposition (HSVD) filter^23^. For Siemens spectra, an additional pre-phasing step was introduced by modelling the signals from creatine and choline-containing compounds at 3.02 and 3.20 ppm with a double Lorentzian model and applying the inverted model phase to the data. This step corrected a zero-order phase shift in the data arising from the HSVD water removal, likely because the Siemens water suppression introduced asymmetry to the residual water signal. Finally, the pre-processed spectra were exported in .RAW format.

### Data modelling

Fully localized 2D density-matrix simulations implemented in the MATLAB toolbox FID-A ^24^ with vendor-specific refocusing pulse information, timings, and phase cycling were used to generate three vendor-specific basis sets (GE, Philips, and Siemens) including 19 spin systems: ascorbate, aspartate, Cr, negative creatine methylene (-CrCH2), γ-aminobutyric acid (GABA), glycerophosphocholine (GPC), glutathione, glutamine (Gln), glutamate (Glu), water (H2O), myoinositol (mI), lactate, NAA, N-acetylaspartylglutamate (NAAG), phosphocholine (PCh), PCr, phosphoethanolamine, scyllo-inositol, and taurine. The -CrCH2 term is a simulated negative creatine methylene singlet at 3.95 ppm, included as a correction term to account for effects of water suppression and relaxation. It is not included in the tCr model, which is used for quantitative referencing.

8 additional Gaussian basis functions were included in the basis set to simulate broad macromolecules and lipid resonances^25^ (simulated as defined in section 11.7 of the LCModel manual^26^): MM0.94, MM1.22, MM1.43, MM1.70, MM2.05, Lip09, Lip13, Lip20. The Gaussian amplitudes were scaled relative to the 3.02 ppm creatine CH3 singlet in each basis set (details in **Supplementary Material 1**). Finally, to standardize the basis set for each algorithm, basis sets were stored as .mat files for use in Osprey and as .BASIS-files for use in LCModel and Tarquin. In the following paragraphs, each LCM algorithm investigated in this study is described briefly (for details, please refer to the original publications^5,7,11^).

### LCModel v6.3

The LCModel (6.3-0D) algorithm^11^ models data in the frequency-domain. First, time-domain data and basis functions are zero-filled by a factor of two. Second, frequency-domain spectra are frequency-referenced by cross-correlating them with a set of delta functions representing the major singlet landmarks of NAA (2.01 ppm), Cr (3.02 ppm), and Cho (3.20 ppm). Third, starting values for phase and linebroadening parameters are estimated by modelling the data with a reduced basis set containing NAA, Cr, PCh, Glu, and mI, with a smooth baseline. Fourth, the final modelling of the data is performed with the full basis set, regularized lineshape model and baseline, with starting values for phase, linebroadening, and lineshape parameters derived from the previous step. Model parameters are determined with a Levenberg-Marquardt^27,28^ non-linear least-squares optimization implementation that allows bounds to be imposed on the parameters. Metabolite amplitude bounds are defined to be non-negative, and determined using a non-negative linear least-squares (NNLS) fit at each iteration of the non-linear optimization. Amplitude ratio constraints on macromolecule and lipid amplitude, as well as selected pairs of metabolite amplitudes (e.g. NAA+NAAG), are defined as in Osprey and Tarquin. The spline baseline is constructed from cubic B-spline basis functions, including one additional knot outside either end of the user-specified fit range, with the number of spline functions being defined by the knot spacing parameter. LCModel constrains the model with three additional regularization terms. Two of these terms penalize a lack of smoothness in the spline baseline and lineshape models using the second derivative operator, preventing unreasonable flexibility of the spline baseline and lineshape irregularity. The third term penalizes deviations of the metabolite Lorentzian linebroadening and frequency shift parameters from their expected values.

### Osprey

The Osprey (1.0.0) frequency-domain LCM algorithm^5^ adopts several key features of the LCModel and Tarquin algorithms. Osprey follows the four-step workflow of LCModel including zero-filling, frequency referencing, preliminary optimization to determine starting values, and final optimization over the real part of the frequency-domain spectrum. The model parameters are zero- and first-order phase correction, global Gaussian linebroadening, individual Lorentzian linebroadening, and individual frequency shifts, which are applied to each basis function. The final model includes the full basis set, as well as unregularized lineshape and spline baseline models. The baseline knot spacing is set to 0.15 ppm for preliminary modelling step with a reduced basis set and increased to 0.4 ppm for the final full model. Similar to LCModel, model parameters are determined with a Levenberg-Marquardt^27,28^ non-linear least-squares optimization algorithm and a NNLS fit to determine the non-negative metabolite amplitudes at each step of the non-linear optimization.

### Tarquin

Tarquin (4.3.10)^7^ uses a four-step approach in the time domain to model spectra. First, residual water is removed using singular value decomposition. Second, the global zero-order phase is determined by minimizing the difference between the magnitude and the real spectra in the frequency domain. Third, zero-filling to double the number of points and frequency referencing are performed, as in the other algorithms. This step also estimates a starting value for the Gaussian linebroadening used in the fourth step, the final modelling. The model includes common Gaussian linebroadening, individual Lorentzian linebroadening, individual frequency-shifts, and zero- and first-order phase correction factors applied in the frequency domain.

Optimization is performed in the time domain with a constrained non-linear least-squares Levenberg-Marquardt solver, allowing bounds and constraints on the parameters. In addition, the range of time-domain datapoints is limited by removing the first 10 ms of the FID, so as to omit the fast-decaying macromolecule and lipid signals. Finally, the baseline is estimated in the frequency domain by convolving the model residual with a Gaussian filter with a width of 100 points.

### Model parameters

The parameters chosen for each tool are summarized in **Figure 1C**. The fit range was limited to 0.5 to 4 ppm in LCModel and Osprey to reduce effects of differences in water suppression techniques. For the baseline handling, the default and most commonly used parameters were chosen, i.e. bLineKnotSpace = 0.4 ppm for Osprey, DKNMNT = 0.15 ppm for LCModel, and an FID range from 10 ms to 50% of the FID for Tarquin.

### Quantification, visualization, and secondary analyses

#### Quantification

The four major metabolite complexes tNAA (NAA + NAAG), tCho (GPC + PCh), mI, and Glx (Glu + Gln) were quantified as basis-function amplitude ratios relative to total creatine (tCr = Cr + PCr). Since the primary purpose was to compare performance of the core LCM algorithms, no additional relaxation correction or partial volume correction was performed.

Model visualizations were generated with the *OspreyOverview* module, which allows LCModel and Tarquin results files (.coord and .txt) to be imported. For each algorithm, the visualization includes site-mean spectra, cohort-mean spectra (i.e. the mean of all spectra), and site- and cohort-mean modelling results (complete model, spline baseline, spline baseline + MM components, and the separate models of the major metabolite complexes).

#### Visualization

As in the default visualizations for the LCModel and Tarquin software interfaces, inverse phase estimates were applied to the spectra and final models. For the visualization, spectra were normalized to the amplitude of the 3-ppm creatine singlet, and a DC offset was added to each site mean spectrum to align the mean frequency-domain amplitude between 1.85 and 4.0 ppm, to aid visual comparison between algorithms and sites.

#### Secondary analyses

To investigate potential vendor differences in linewidth and SNR based on the different export formats of the data, mean and standard deviation of the NAA linewidth and SNR were investigated.

To investigate baseline power variability unbiased by DC offsets, the MM + baseline models were first aligned vertically according to the frequency-domain minimum of the acquired spectra between 2.66 and 2.7 ppm (i.e. between the aspartyl signals, which is the region with the highest consistency between the baseline models). Baseline models were normalized to the frequency-domain amplitude of each metabolite spectrum between 2.9 and 3.1 ppm to account for differences in the scaling of the model outputs of LCModel and Tarquin. Baseline power within the frequency ranges was then defined as the range-normalized integral of the baseline model between 0.5 and 1.95 ppm; 1.95 and 3.6 ppm and 3.6 to 4.0. The baseline power variability was then defined as the standard deviation of the baseline power calculated across all subjects.

To similarly investigate potential interactions between baseline power and metabolite estimates, the baseline power beneath each major metabolite was then defined as the range-normalized integral of the baseline model between 1.9 and 2.1 ppm for the tNAA baseline; 3.1 and 3.3 ppm for the tCho baseline; 3.33 and 3.75 ppm for mI; and 1.9 to 2.5 ppm and 3.6 to 3.8 ppm for the Glx baseline.

The contribution of variance in modelling of the creatine reference signal to metabolite ratios was also investigated. To this end, each individual total creatine model (Cr + PCr) was normalized to the frequency-domain amplitude of each metabolite spectrum between 1.9 and 2.1 ppm to account for differences in the scaling of the total creatine model outputs of LCModel and Tarquin. Finally, the integral over the individual creatine model was calculated.

Additionally, water referenced tCr concentrations were calculated as the ratio of the tCr and water amplitude of each algorithm. No further corrections were applied to these tCr estimates.

#### Data analysis

Quantitative metabolite estimates (tNAA/tCr, tCho/tCr, mI/tCr, Glx/tCr) were statistically analyzed and visualized using R^29^ (Version 3.6.1) in RStudio (Version 1.2.5019, RStudio Inc.). The functions are publicly available^30^. The supplemental materials with MATLAB- and R-files, example LCModel control files (one for each vendor), and Tarquin batch-files for this study are publicly available^31^. The results from each LCM algorithm were imported into R with the *spant* package^32^.

#### Distribution analysis

The results are presented as raincloud plots^33^ and Pearson’s correlation analysis using the *ggplot2* package^34^. The raincloud plots include individual data points, boxplots with median and 25^th^/75^th^ percentiles, a smoothed distribution, and mean ± SD error bars to identify systematic differences between the LCM algorithms. In addition, the coefficient of variation (CV = SD/mean) and the mean 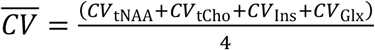 across all four metabolites of each algorithm are calculated.

#### Correlation analysis

The Pearson’s correlation analysis featured different levels, including pair-wise correlations between algorithms, as well as correlations between baseline power and metabolite estimates of each algorithm. The pair-wise coefficient of determination on the global level (black R^2^), as well as within-vendor coefficient of determination (color-coded R^2^) with different color shades for different sites are reported. Furthermore, mean 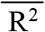 for each pair-wise coefficient of determination (e.g. Osprey vs LCModel) and metabolite, estimated by row or column means e.g.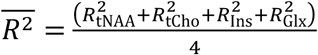, and a cohort mean 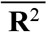 (across all pair-wise correlations) are calculated. The correlations were Bonferroni corrected for the number of correlation tests. The cohort mean 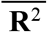 was used to identify global associations across all correlation analysis, while the mean 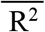 allowed the identification of algorithm-specific (row means) and metabolite-specific (column means) interactions across all correlation analysis. Associations between the outcome of specific algorithms were identified by the pair-wise correlation analysis (R^2^). Vendor-specific effects were identified by differentiating between global level and within-vendor correlations.

#### Statistical analysis

In the statistical analysis, the presence of significant differences in the mean and the variance of the metabolite estimates was assessed. Global metabolite estimates were compared between algorithms with parametric tests, following recommendations for large sample sizes^35^. The data were not grouped by vendor or site, and the statistical tests were set up as paired without any further inference. Differences of variances were tested with Fligner-Killeen’s test with a post-hoc pair-wise Fligner-Killeen’s test and Bonferroni correction for the number of pair-wise comparisons. Depending on whether variances were different or not, an ANOVA or Welch’s ANOVA was used to compare means with a post-hoc paired t-test with equal or non-equal variances, respectively.

#### Linear mixed-effects analysis

Linear mixed-effects models were set up as a repeated-measure analysis to determine variance partition coefficients to assess the contributions of algorithm-, vendor-, site-, and participant-specific effects to the total variance. A log-likelihood statistic was used to calculate the goodness of fit. The probability of observing the test statistic was evaluated against the null hypotheses, which was simulated by performing parametric bootstrapping (2000 simulations) ^36^.

## Results

All 277 spectra were successfully processed, exported, and quantified with the three LCM algorithms; no modelled spectra were excluded from further analysis.

### Summary and visual inspection of the modelling results

**Figure 2** shows the 277 spectra, models and residuals for each algorithm (A-C) color-coded by vendor. In general, the phased spectra and models agreed well between vendors for all algorithms. The most notable differences in spectral features are visible between 0.5 and 1.95 ppm with each algorithm modelling the macromolecules and apparent fat peaks as baseline or macro-molecule basis function to a different degree. Similarly, the baseline between 3.6 and 4 ppm is estimated differently by each algorithm, which is changing the amplitude of the residual in this frequency range and potentially the metabolite estimates. Comparing the algorithms, notable differences in spectral features and inter-subject variability in the estimated baseline models appeared between 0.5 and 1.95 ppm (SD baseline area: 0.45 (Osprey) > 0.36 (Tarquin) > 0.21 (LCModel)) and between 3.6 and 4 ppm (SD baseline area: 0.15 (LCModel) > 0.13 (Osprey) > 0.10 (Tarquin)) (as shown in **Figure 2A-C** and calculated in the secondary analysis). A high agreement in the estimated baseline models was found between 1.95 and 3.6 ppm (SD baseline area: 0.04 (Osprey) > 0.03 (LCModel) = 0.03 (Tarquin)).

**Figure 2.**
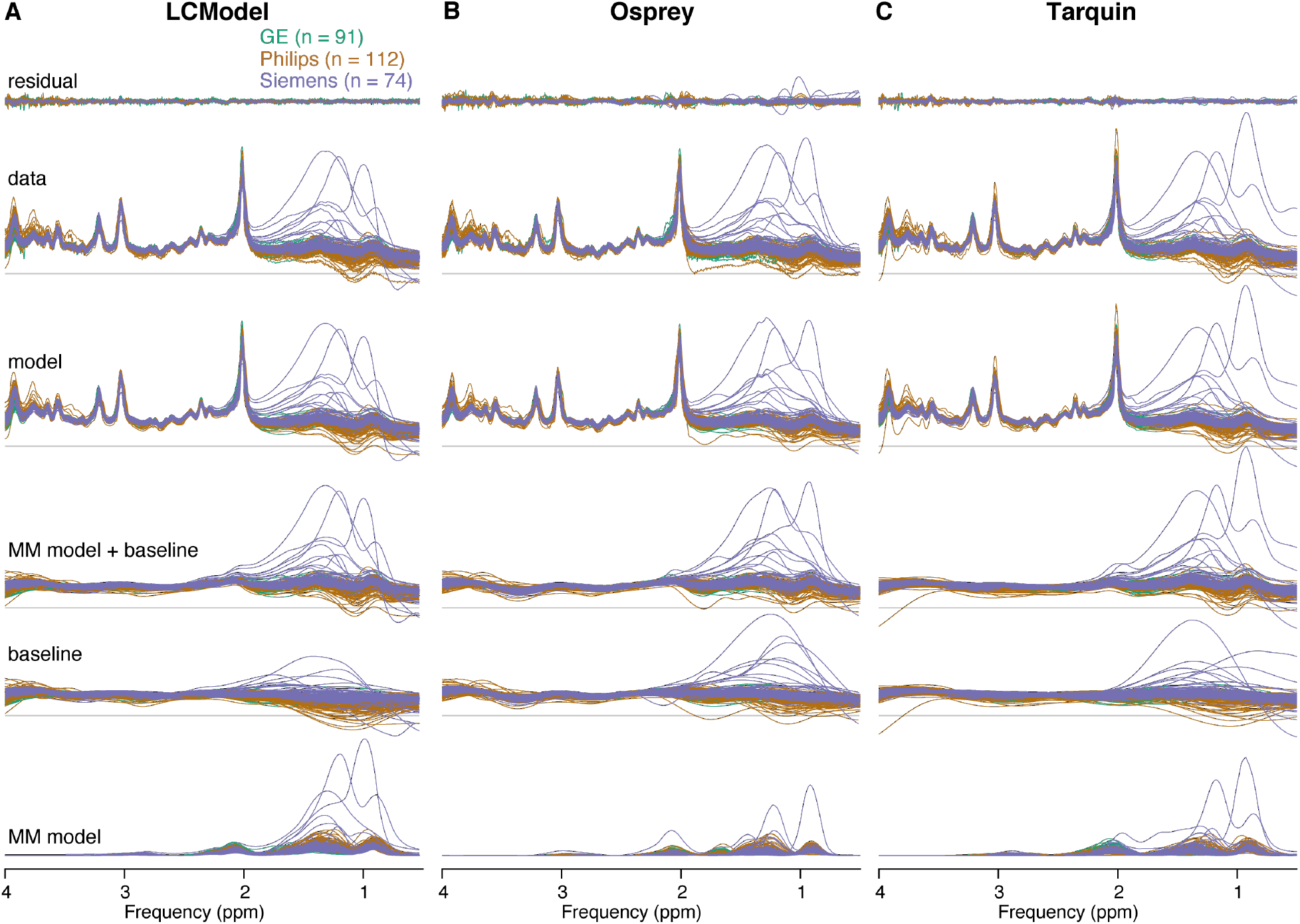
Summary of the individual modelling results. (A–C) individual residuals, data, models, MM models + baseline, baseline and MM models for each LCM algorithm, color-coded by vendor.

A site-level averaged summary is shown in **Figure 3A, B** and **C**, for analyses in LCModel, Osprey, and Tarquin, respectively. The averaged data, models and residuals for each of the 25 sites are color-coded by vendor. The cohort-mean of all analyses for each vendor is shown in **Figure 3D, E** and **F** (GE, Philips and Siemens, respectively). Data, models and residuals are color-coded by algorithm.

**Figure 3.**
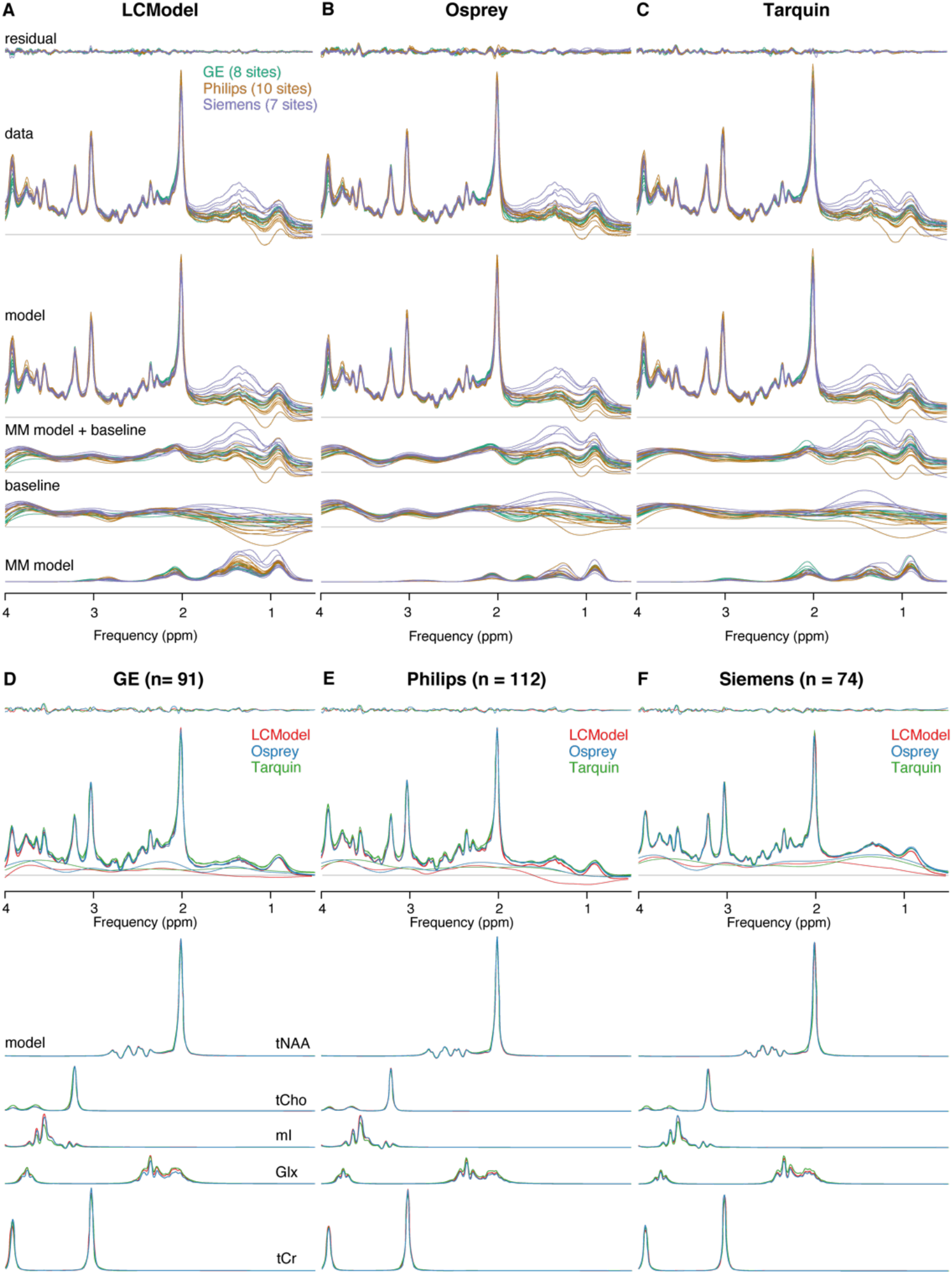
Summary of the modelling results. (A–C) site-level averaged residual, data, model, MM model + baseline, baseline and MM model for each LCM algorithm, color-coded by vendor. (D–F) cohort-mean residual, data, model, MM model + baseline, and metabolite models for each vendor, color-coded by LCM algorithm.

Cohort-mean spectra and models agreed well across all vendors and algorithms (**Figure 3D-F**). The greatest differences in the spectral features of the baseline between algorithms occur between 0.5 and 1.95 ppm, with closer agreement between Osprey and Tarquin than with LCModel. The amplitude of the residual over the whole spectral range is highest for Osprey, and similar for Tarquin and LCModel.

The mean NAA linewidth was significantly lower (p < 0.001) for Philips (6.3 ± 1.3 Hz) compared to GE (7.3 ± 1.5 Hz), while no differences in the linewidth were found for the other comparisons (Siemens 6.6 ± 2.4 Hz). The mean SNR was significantly higher for Siemens (285 ± 72) compared to both other vendors (p < 0.001) and significantly higher (p < 0.001) for Philips (226 ± 58) compared to GE (154 ± 37).

### Metabolite level distribution

The tCr ratio estimates and CVs of the four metabolites are summarized in **Table 2**. Distributions and group statistics are visualized in **Figure 4**, with the four rows corresponding the three vendors and a cohort summary across all datasets.

**Table 1.** Metabolite level distribution. Mean, standard deviation and coefficient of variation (CV) of each metabolite-to-creatine ratio, listed by algorithm and vendor as well as global summary values. Asterisks indicate significant differences (adjusted p < 0.01 = ** and adjusted p < 0.001 = *** or ^###^ or ’’’) in the mean (for the metabolite ratios) or the variance (for the CV) compared to the algorithm in the next row (LCModel vs Osprey = ** or ***, Osprey vs Tarquin = ^###^, and Tarquin vs LCModel = ’’’).

| | [metabolite] / [tCr] (mean ± SD) | | | |
| --- | --- | --- | --- | --- |
|  | tNAA | tCho | mI | Glx |
| **GE** |  |  |  |  |
| LCModel | 1.48 ± 0.12 | 0.19 ± 0.02 | 0.85 ± 0.10 | 1.75 ± 0.25 |
| Osprey | 1.47 ± 0.10 | 0.18 ± 0.02 | 0.78 ± 0.09 | 1.42 ± 0.17 |
| Tarquin | 1.48 ± 0.11 | 0.22 ± 0.03 | 0.57 ± 0.07 | 2.05 ± 0.22 |
| **Philips** |  |  |  |  |
| LCModel | 1.38 ± 0.10 | 0.17 ± 0.02 | 0.81 ± 0.08 | 1.46 ± 0.14 |
| Osprey | 1.50 ± 0.12 | 0.18 ± 0.02 | 0.86 ± 0.10 | 1.34 ± 0.16 |
| Tarquin | 1.40 ± 0.12 | 0.16 ± 0.03 | 0.60 ± 0.09 | 1.78 ± 0.19 |
| **Siemens** |  |  |  |  |
| LCModel | 1.52 ± 0.19 | 0.19 ± 0.02 | 0.83 ± 0.09 | 1.65 ± 0.31 |
| Osprey | 1.54 ± 0.12 | 0.19 ± 0.02 | 0.89 ± 0.06 | 1.45 ± 0.14 |
| Tarquin | 1.50 ± 0.15 | 0.18 ± 0.03 | 0.65 ± 0.07 | 2.04 ± 0.19 |
| **global** |  |  |  |  |
| LCModel | 1.45 ± 0.15*** | 0.18 ± 0.02 | 0.83 ± 0.09 | 1.45 ± 0.15*** |
| Osprey | 1.50 ± 0.12### | 0.18 ± 0.02 | 0.84 ± 0.09### | 1.50 ± 0.12### |
| Tarquin | 1.46 ± 0.14 | 0.18 ± 0.04 | 0.60 ± 0.08’’ | 1.93 ± 0.24’’ |
|  | CV (SD/mean) |  |  |  |
|  | tNAA | tCho | mI | Glx |
| **GE** |  |  |  |  |
| LCModel | 7.9% | 12.9% | 11.8% | 14.2% |
| Osprey | 6.9% | 9.7% | 11.1% | 11.8% |
| Tarquin | 7.5% | 11.7% | 11.2% | 10.8% |
| **Philips** |  |  |  |  |
| LCModel | 7.2% | 10.6% | 9.9% | 9.7% |
| Osprey | 8.0% | 10.0% | 11.8% | 11.9% |
| Tarquin | 8.8% | 19.8% | 15.2% | 10.7% |
| **Siemens** |  |  |  |  |
| LCModel | 12.4% | 13.4% | 10.8% | 18.7% |
| Osprey | 8.0% | 11.1% | 6.9% | 10.0% |
| Tarquin | 10.1% | 14.3% | 10.5% | 9.3% |
| **global** |  |  |  |  |
| LCModel | 10.0% | 13.2%** | 10.9% | 16.4%*** |
| Osprey | 7.8% | 10.4%### | 11.7%### | 11.8%### |
| Tarquin | 9.3% | 20.5%’’ | 13.6% | 12.3% |

**Figure 4.**
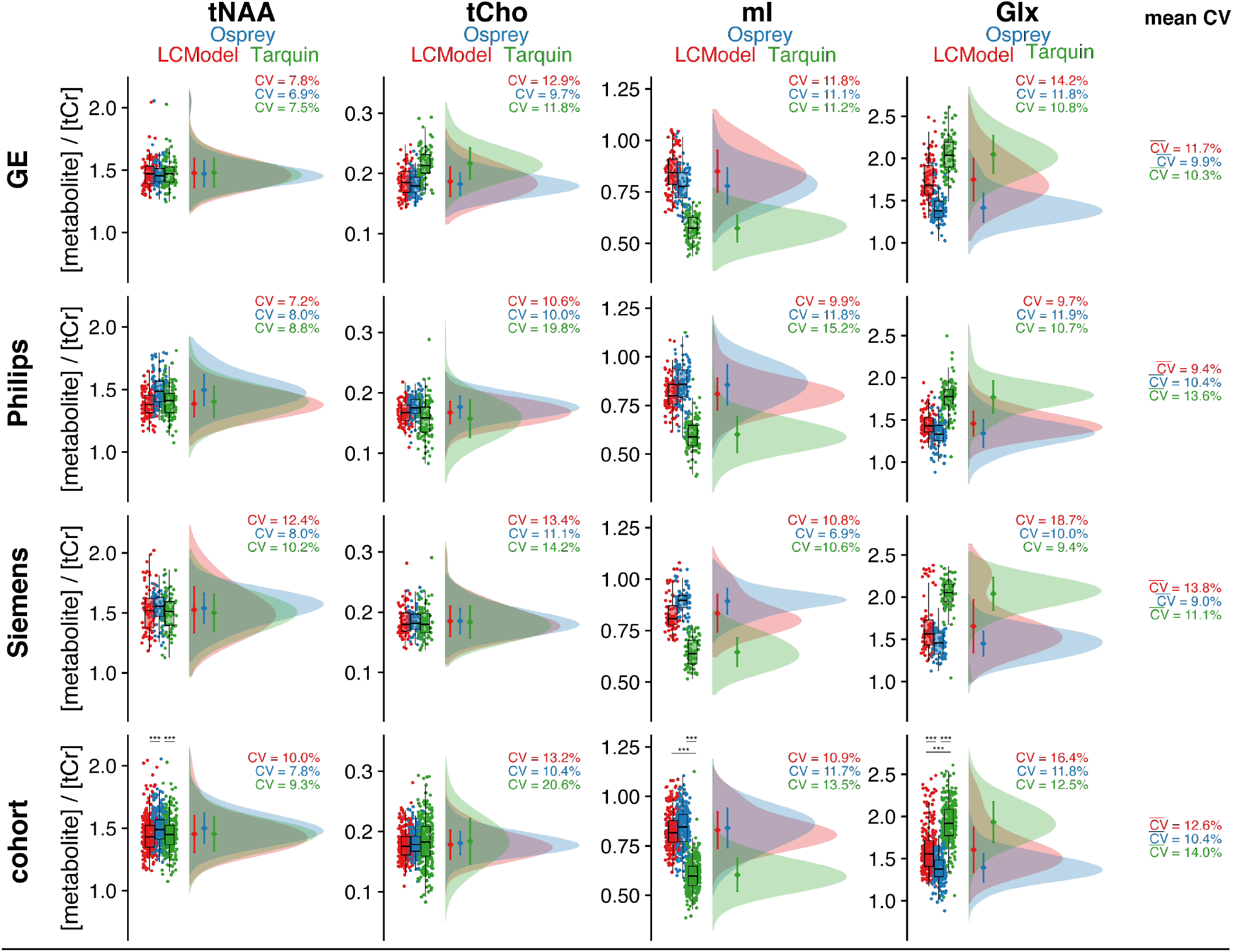
Metabolite level distribution. Raincloud plots of the metabolite estimates of each LCM algorithm (color-coded). The four metabolites are reported in the columns, and the three vendors in rows, with a cohort summary in the last row. The coefficient of variation is reported for each distribution, as well as a mean 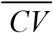 reported in the last column, which is calculated across each row. Asterisks indicate significant differences (adjusted p < 0.001 = ***).

Between-algorithm agreement was greatest for the group means and CVs of tNAA and tCho. The cohort-mean CV was lowest for Osprey (10.4%), followed by LCModel (12.6%) and Tarquin (14.0%). Group means and CVs for tNAA are relatively consistent. As a result, the cohort-mean tNAA/tCr was 1.45 ± 0.15 for LCModel, 1.50 ± 0.12 for Osprey, and 1.45 ± 0.14 for Tarquin, with significant differences between Osprey and both other LCM algorithms.

Cohort means for tCho showed a high agreement between all algorithms. The global CV of tCho estimates was significantly higher for Tarquin compared to both other algorithms, and significantly lower for Osprey compared to LCModel. Global tCho/tCr was 0.18 ± 0.02 for LCModel, 0.18 ± 0.02 for Osprey, and 0.18 ± 0.04 for Tarquin.

For mI, group means and CVs were comparable for Osprey and LCModel, while Tarquin estimates were lower by about 25%. Global CVs were significantly lower for Osprey compared to Tarquin, while no significant differences in the CV were found for the other comparisons. Global mI/tCr was 0.83 ± 0.09 for LCModel, 0.84 ± 0.09 for Osprey, and 0.60 ± 0.08 for Tarquin, with significant mean differences between all Tarquin and both other algorithms.

Group means and CVs for Glx were comparable between Osprey and LCModel, while estimates were about 30% higher in Tarquin. Global CV was significantly lower for Osprey compared to both other algorithms. Global Glx/tCr was 1.45 ± 0.15 for LCModel, 1.50 ± 0.12 for Osprey, and 1.93 ± 0.24 for Tarquin, with significant differences between all algorithms. Mean 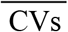, estimated by the row-mean, were between 9.0 and 13.8% for all algorithms and vendors.

### Correlation analysis: pairwise comparison between LCM algorithms

The correlation analysis for each metabolite and algorithm pair is summarized in **Figure 5**. 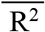 for each algorithm pair and metabolite are reported in the corresponding row and column, respectively.

**Figure 5.**
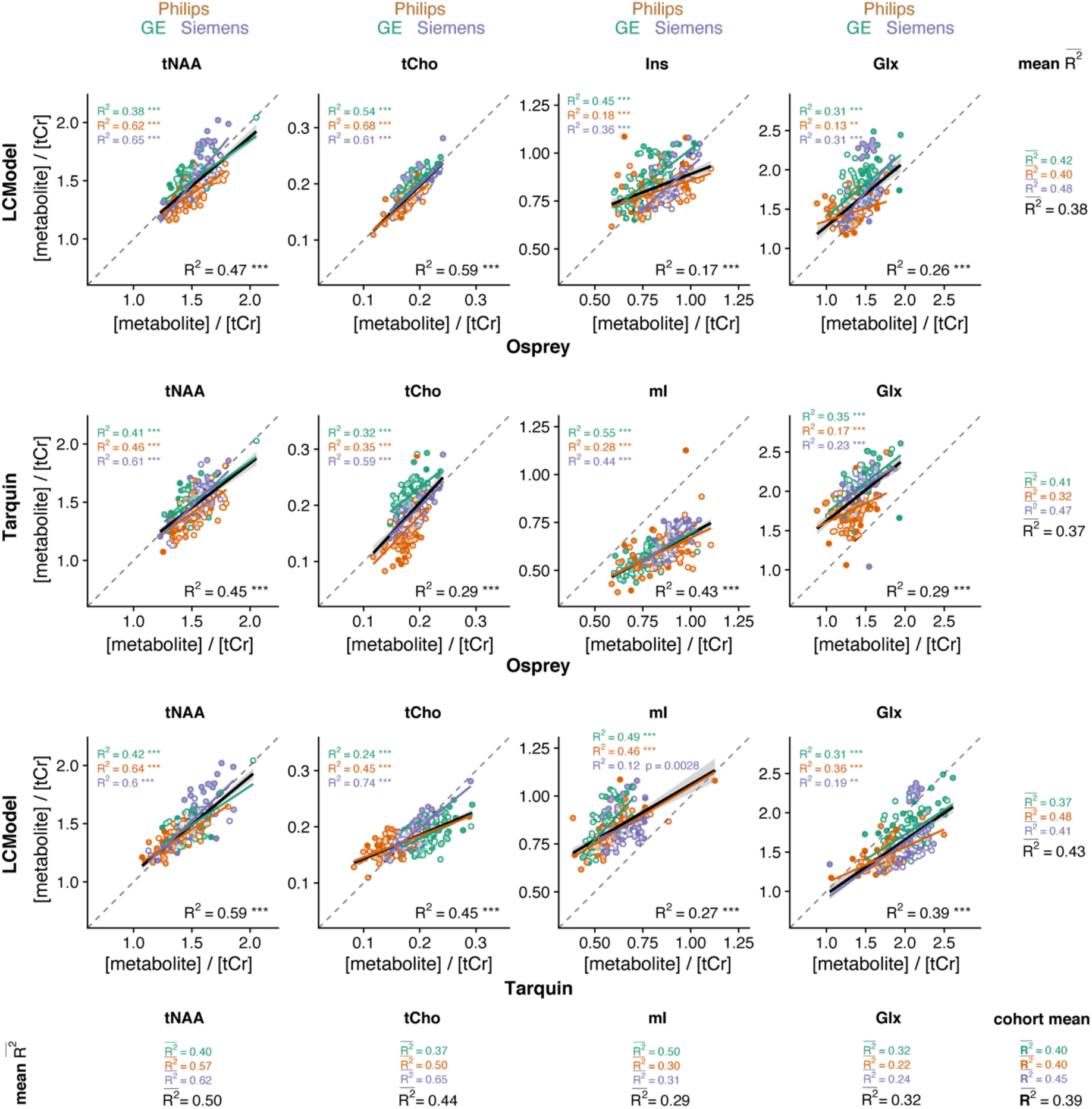
Pairwise correlational comparison of algorithms. LCModel and Osprey are compared in the first row, Tarquin and Osprey in the second row, and LCModel and Tarquin in the third row. Each column corresponds to a different metabolite. Within-vendor correlations are color-coded; global correlations are shown in black. The 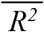 values are calculated along each dimension of the grid with mean R^2^ for each metabolite and each correlation. A cohort-mean 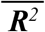 value is also calculated across all twelve pair-wise correlations. Asterisks indicate significant correlations (adjusted p < 0.01 = ** and adjusted p < 0.001 = ***).

The cohort-mean 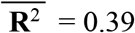 suggests an overall moderate agreement between metabolite estimates from different algorithms. The agreement between algorithms, estimated by the row-mean 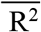, was highest for Tarquin-vs-LCModel 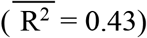, followed by Osprey-vs-LCModel 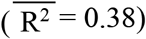, and Osprey-vs-Tarquin 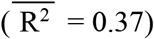.

The agreement between algorithm for each metabolite, estimated by the column-mean 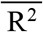, was highest for tNAA 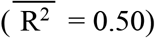, followed by tCho 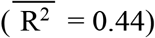, Glx 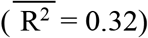, and mI 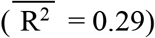. The cohort-mean 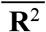 for each vendor was higher for Siemens 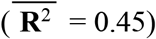 than for GE 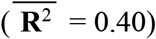 and Philips 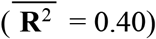.

While the within-metabolite mean 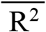 (average down the columns in **Figure 5**) are comparable between vendors, there is substantially higher variability of the R^2^ values with increasing granularity of the analysis. **Supplementary Material 2** includes an additional layer of correlations at the site level.

### Correlation analysis: baseline and metabolite estimates

The correlation analysis between local baseline power and metabolite estimates for each algorithm is summarized in **Figure 6**. The cohort-mean 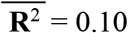 suggests that overall, there is an association between local baseline power and metabolite estimates, that is weak but statistically significant. The influence of baseline on metabolite estimates differs between metabolites, as reflected by the column-mean 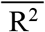 which was lowest for tCho 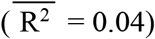 and tNAA 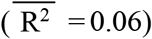, and higher for mI 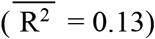 and Glx 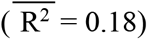. The global baseline correlations all had negative slope, except for tCho estimates of Tarquin.

**Figure 6.**
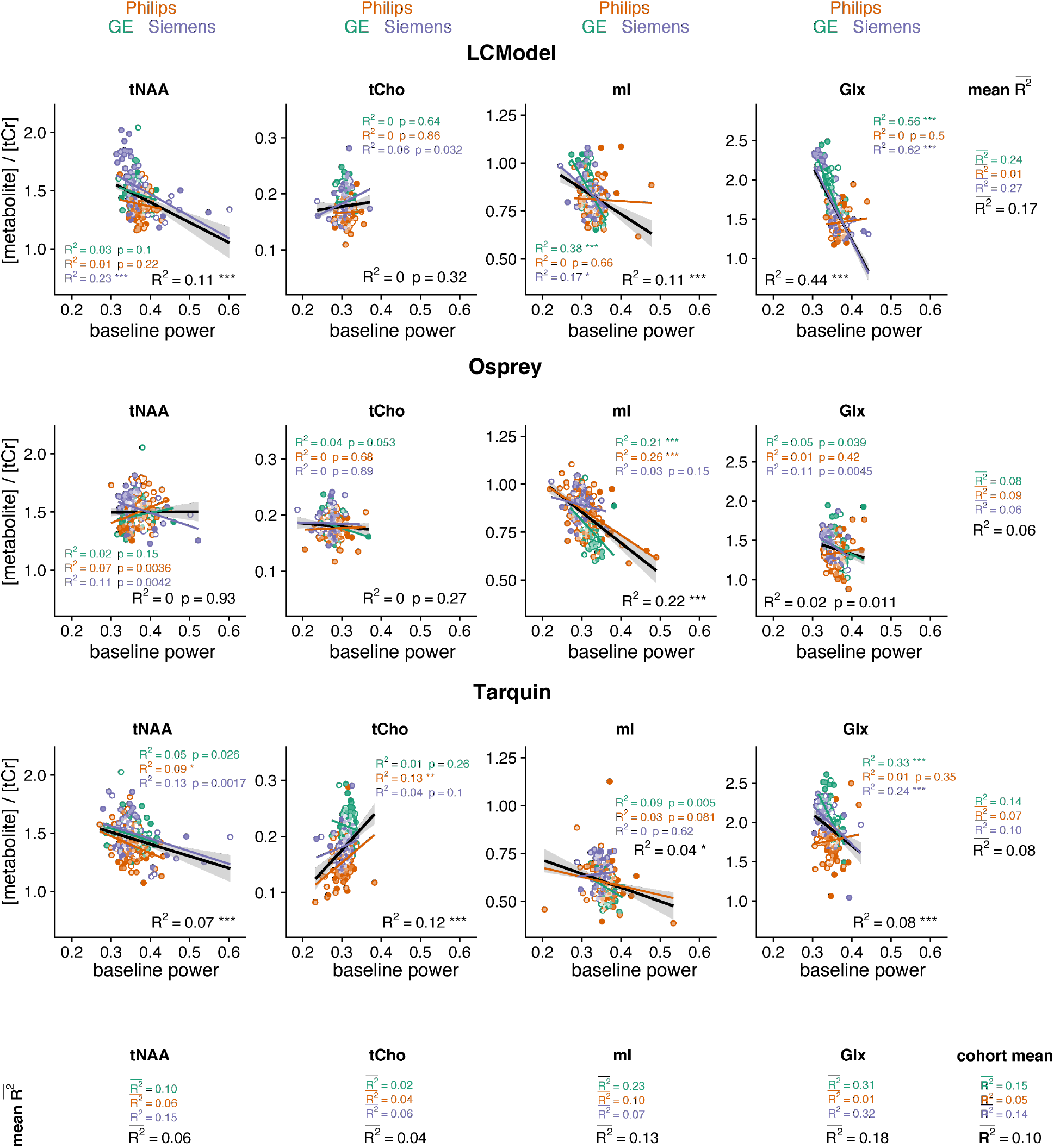
Correlation analysis between metabolite estimates and local baseline power for each algorithm, including global (black) and within-vendor (color-coded) correlations. The mean 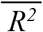 values are calculated along each dimension of the grid for each metabolite and each algorithm. Similarly, a cohort-mean 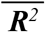 value is calculated across all twelve pair-wise correlations. Asterisks indicate significant correlations (adjusted p < 0.05 = *, adjusted p < 0.01 = **, adjusted p < 0.001 = ***).

The mean 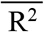 across metabolites for each algorithm, calculated as the row mean, were low for all algorithms with LCModel 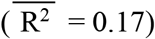 showing a greater effect than Tarquin 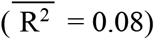 and Osprey 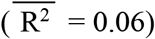. Comparing between vendors, the cohort-mean 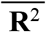 was higher for GE 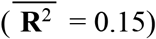 and Siemens 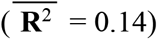 than for Philips 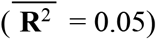 spectra.

### Variability of total creatine models

Mean tCr model spectra (± one standard deviation) are summarized in **Supplementary Material 3A** for each vendor and LCM algorithm, along with distribution plots of the area under the model.

The agreement in mean and CV is greatest between Osprey and Tarquin for all vendors, while tCr areas for LCModel appear slightly higher. Differences in water suppression are accounted for with the -CrCH2 correction term, which is not included in the tCr model used for quantitative referencing.

**Supplementary Material 3B** shows the distribution of the water referenced tCr concentrations with a high agreement of CV between all algorithms and vendors. The agreement between the mean was higher between Osprey and Tarquin for GE and Siemens, while the mean concentrations were different for LCModel. The highest variation between algorithms was found for Philips.

### Linear mixed-effect models

The results from the linear mixed-effects model analysis are summarized in Table 3. The algorithm-specific effect ranged between 0.7% (tCho) and 58.7% (mI) and was significant for all metabolites. Significant vendor-specific effects were found for Glx (10.1%) and tCho (17.5%), while significant site-specific effects ranged between 3.8% (mI) and 21.7% (tNAA). The participant-specific effects ranged from 7.5 (Glx) to 40.4% (tNAA) and was significant for all metabolites. The metabolite distribution divided by algorithm, vendor, site, and subject is shown in **Figure 7**.

**Table 3.**
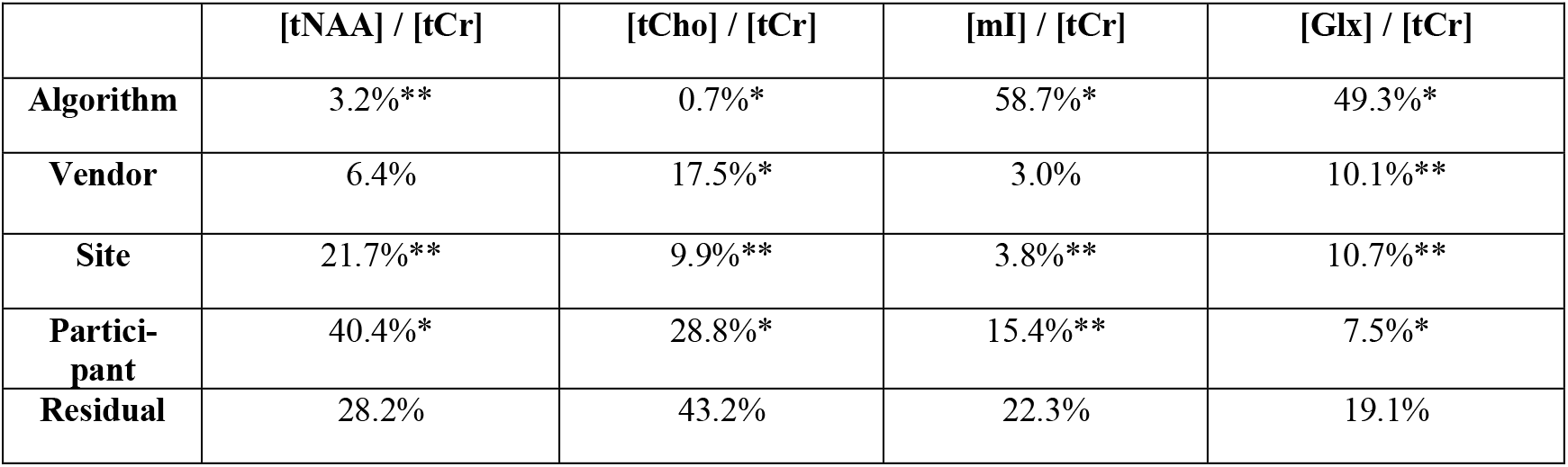
Variance partition coefficients for algorithm-, vendor, site-, and participant-level effects for the metabolite levels (shown as percentage). The residual represents the part of the total variance which is not explained by the linear mixed-effect model. Asterisks indicate significant effects based on linear mixed-effect modelling (p < 0.05 = * and p < 0.01 = **).

**Figure 7.**
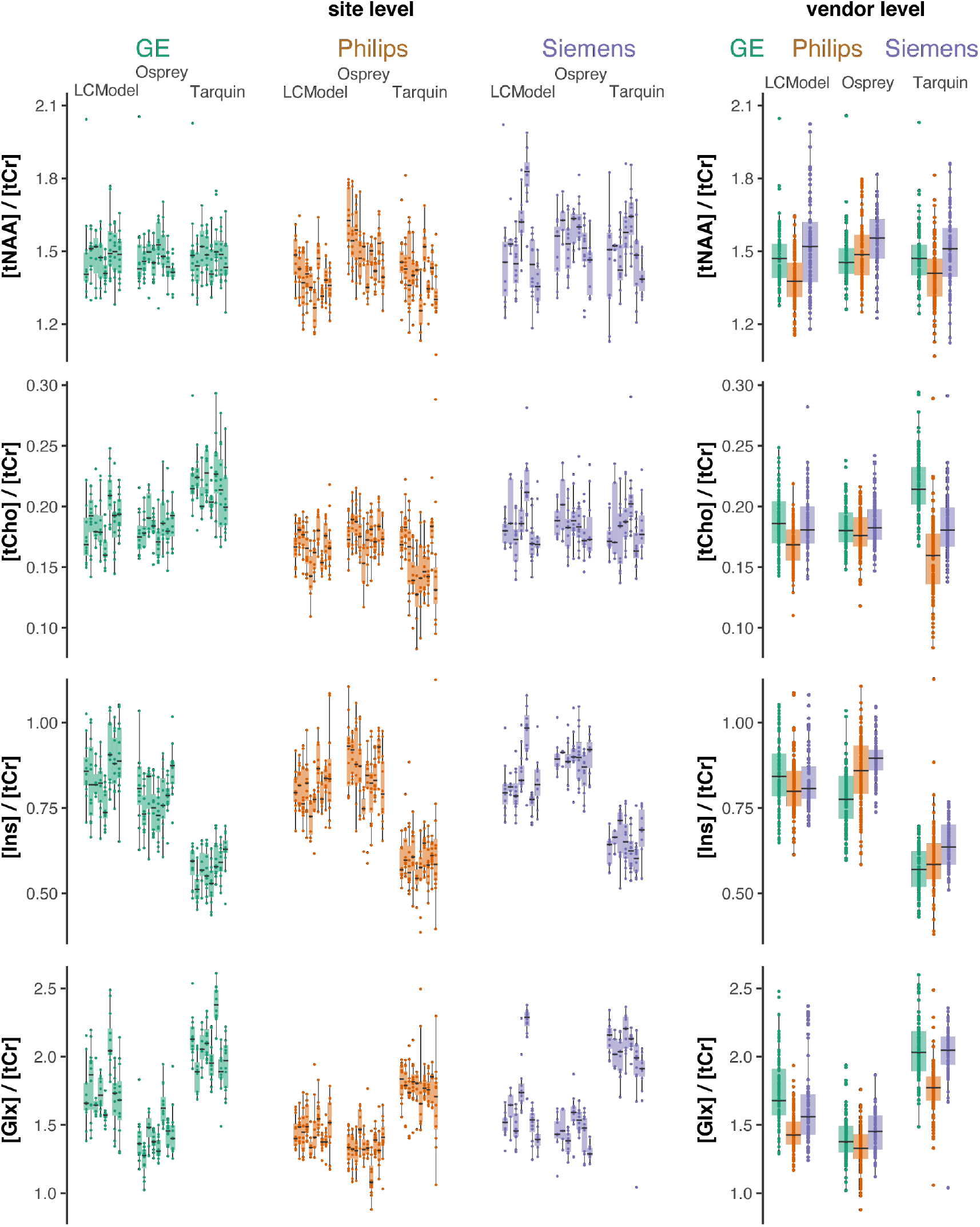
Metabolite level distribution by site. Boxplots of the metabolite estimates of each LCM algorithm and vendor (color-coded). Each site is represented as a single box with individual data points The four metabolites are reported in the rows and the three vendors are represented in the columns, Column four represents the vendor collapsed distributions.

## Discussion

We have presented a three-way comparison of LCM algorithms applied to a large dataset of short-TE in-vivo human brain spectra. The aims at the onset were to compare metabolite estimates obtained with different LCM algorithms, as applied in the literature, and to identify potential sources of differences between the algorithms. The major findings are:

- Group means and CVs for tNAA and tCho agreed well across vendors and algorithms. For mI and Glx, group means and CVs were less consistent between algorithms, with a higher degree of agreement between Osprey and LCModel than with Tarquin.
- The strength of the correlations between individual metabolite estimates from different algorithms was moderate. In general, tNAA and tCho estimates from different algorithms agreed better than Glx and mI. With each sub-level of analysis, the variability of correlation strength increased, i.e. correlations grew increasingly variable when calculated separately for each vendor, or even each site.
- Overall, the association between metabolite estimates and the local baseline power was significant, with mI and Glx showing stronger associations than tNAA and tCho, and LCModel showing greater effects than Tarquin and Osprey.
- A large vendor-specific effect was found for Glx (49.3%) and mI (58.7%) by calculating variance partition coefficients using linear mixed-effects modelling. For tNAA and tCho, the participant-level effect was largest with 40.4% and 28.8%, respectively.

The strong agreement of group means and CVs for metabolites with prominent singlets (tNAA/tCho) and inconsistency for lower-intensity coupled signals (mI/Glx) are in line with previous two-tool comparisons of simulated data ^7,15^ and in-vivo studies with smaller sample sizes 7,14,16.

While previous work highlighted group means and standard deviations, the between-algorithm agreement of individual metabolite estimates has not been extensively studied. Our results suggest that substantial variability is introduced by the choice of the analysis software itself, indicated by only moderate between-algorithm correlation strength (between-algorithm mean 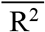 <= 0.5 for all investigated metabolites), even for the well-established LCM algorithms LCModel and Tarquin (R^2^ between 0.27 and 0.59 for all metabolites). This finding raises concerns about the generalizability and reproducibility of MRS study results. MRS studies typically suffer from low sample sizes (∼20 per comparison group is common). Considering the moderate between-tool correlation of individual estimates, it is likely that marginally significant group effects and correlations found with one analysis tool will not be found with another tool, even if the exact same dataset is used. This is exacerbated by the substantial variability of correlation strengths at vendor- or even site-level, and is even more likely to be the case for ‘real-life’ clinical data, given the relatively high quality of the dataset ‘best case scenario’ in this study (standardized pre-processing; large sample size; high SNR; low linewidth; young, healthy, cooperative subjects). While two previous studies found that some differences between clinical groups remained significant independent of the LCM algorithm ^14,16^, this is questionable as a default assumption. The lack of comparability arising from the additional variability originating in the choice of analysis tool is rarely recognized or acknowledged. If choice of analysis tool is a significant contributor to measurement variance, it could be argued that modelling of data with more than one algorithm will improve the robustness and power of MRS studies. It should also be investigated whether the reduction of the degrees of freedom by improving MM and baseline models (e.g. by using acquired MM data) increases between-tool agreement and consistency between sites and vendors.

### Sources of variance

In order to understand the substantial variability introduced by the choice of analysis tool, the influence of modelling strategies and parameters on quantitative results needs to be better understood. Previous investigations have shown that, within a given LCM algorithm, metabolite estimates can be affected by the choice of baseline knot spacing^37,38^, the modelling of MM and lipids 37,39, and SNR and linewidth^40–43^. In this study, we focused on the comparison of each LCM with their default and commonly used parameters, and observed differences resulting both from the default parameters and from differences in the core algorithm. Minor differences in spectral quality (SNR and LW) were found between vendors. The agreement between vendors was high for the mean metabolite levels and the cohort-mean correlations. Further vendor-specific effects on the LCM estimation of this dataset are described elsewhere^17^.

LCM relies on the assumption that broad background and baseline signals can be separated from narrower metabolite signals. This is true to a limited degree, and the choice of MM and baseline modelling influences the quantification of metabolite resonances^4^. Our secondary analysis of the relationship between baseline power and metabolite estimates showed a stronger interaction for the broader coupled signals of Glx and mI than the singlets. tCho showed the weakest effect, and the three LCMs showed the highest agreement between the MM+baseline models around 3.2 ppm. The higher variance of Glx and mI estimates may at least partly be explained by the absence of MM basis functions for frequencies >3 ppm in the model. MM signal must therefore either be modelled by metabolite basis functions or the spline baseline. This also emphasizes that any LCM implementation requires a baseline model to account for broad background and baseline signals, with the disadvantage of this being a main source of variance between algorithms. This is also implied by the large algorithm-level effect for mI and Glx in the linear mixed-effects model. Comparing the variance patriation coefficients of this study with the prior single LCModel analysis ^17^ shows a high agreement in the variance partition for tNAA and tCho. A lower agreement is found for mI and Glx as the estimates, of those metabolites strongly differ between algorithms, introducing a high algorithm-level effect. Including experimental MM acquisitions into studies may reduce the degrees of freedom of modelling, but introduce other sources of variance, such as age-dependency^44^ or tissue composition^39,45^. While consensus is emerging that such approaches are recommended many open questions must be resolved before the recommendations can be broadly implemented^25^ and the default LCModel MM basis functions are still commonly used in studies applying MRS. A literature review of the thirty most recent LCModel^11^ citations (Google Scholar, November 10, 2020) in short-TE 3T MRS application studies revealed that 24 of these studies used the default MM basis functions, 1 was performed without MM basis functions at all, and 1 employed a measured MM spectrum. The remaining 4 studies did not report sufficient details, and authors did not respond to our inquiries.

For all three LCM algorithms, optimization between the model and the data is solved by local optimization. Algorithms could converge on a local minimum, if the search space of the non-linear parameters is of high dimensionality, or if the starting values of the parameters are far away from the global optimum^46^. The availability of open-source LCM such as Tarquin and Osprey will allow further investigation of the relationship between optimization starting values and modelling outcomes.

The inherent differences between frequency-domain modelling, which is normally restricted to a specific frequency range, and time-domain modelling, which includes the full spectrum, is a potential source of variance, as the approaches differ in their susceptibility to the residual water peak. In this study, Tarquin’s internal default SVD water removal was included in addition to the HSVD filter in Osprey’s processing pipeline to reduce this effect and to follow Tarquin’s default approach. A secondary analysis confirmed that the effect of Tarquin’s SVD on the metabolite estimates was negligible ^31^. Further standardization between Tarquin and the frequency-domain modelling approaches could be achieved by restricting the frequency range of the basis set accordingly.

Since this study focused on reporting tCr ratios, it is important to consider the variance of the creatine model of each algorithm. With MRS only quantitative in a relative sense, separating the variance contribution of the reference signal is a challenge. While mean tCr model areas were slightly higher for LCModel than for Osprey and Tarquin, there was no generalizable observation of lower tCr ratios from LCModel. CVs of the tCr model areas were comparable across LCM algorithms for each vendor. This is also the case for the water referenced tCr concentrations, which showed no systematic differences in the mean or CV between algorithms. Vendor differences in the water suppression were minimized by limiting the analysis range to 0.5 to 4 ppm, and by including a -CrCH2 correction term (omitted from calculations of the tCr ratios and the secondary analysis of the tCr models). The contribution of the reference signal to the variance of metabolite estimates is unclear and hard to isolate. Nevertheless, tCr referencing was preferred in this study, since water referencing is likely to add additional tool-specific variance resulting from water amplitude estimation.

### Limitations

As mentioned in greater detail above, there is currently no widely adopted consensus on the definition of MM basis functions, and measured MM background data are not widely available to non-expert users. To reflect common practice in current MRS applications, the default MM basis function definitions from LCModel were adapted for each algorithm in this study. These basis functions only included MMs for frequencies < 3.0 ppm, which is likely insufficient for the modelling of MM signals between 3 and 4 ppm^47^, and will have repercussions for the estimation of tCho, mI, and Glx. Second, standard modelling parameters were chosen for each LCM, which ensure a broader comparability to the current literature, but may not be ideal. Third, there is obviously no ‘gold standard’ of metabolite level estimation to validate MRS results against. The performance of an algorithm is often judged based on the level of variance, but low variance clearly does not reflect accuracy and may indicate insufficient responsiveness of a model to the data. Therefore, while comparing multiple algorithms, a higher degree of correlation in the results does not necessarily imply a higher reliability, but it could equally be the case that shared algorithm-based sources of variance increase such correlations. Efforts to use simulated spectra as a gold-standard, including those applying machine learning ^48,49^, can only be successful to the extent that simulated data are truly representative of in-vivo data. Fourth, another criterion to judge the performance of an algorithm is the residual. For example, a small residual indicates a higher agreement between the complete model and the data for LCModel, it does not imply a better estimation of individual metabolites, and may result from the higher degree of freedom in the baseline of LCModel (higher number of splines) compared to Osprey and Tarquin. This is emphasized by the high agreement of the mean mI models, but lower agreement of the baseline models around 3.58 ppm between LCModel and Osprey. Fifth, this study was limited to the two most widely used algorithms LCModel and Tarquin, as well as the Osprey algorithm that is under ongoing development in our group. While including additional algorithms would increase the general understanding of different algorithms, the complexity of the resulting analysis and interpretation would be overwhelming and beyond the scope of a single publication.

## Conclusion

This study presents a comparison of three LCM algorithms applied to a large-scale multi-site short-TE PRESS multi-vendor dataset. While different LCM algorithms’ estimates of major metabolite levels agree broadly at a group level, correlations between results are only weak-to-moderate, despite standardized pre-processing, a large sample of young, healthy and cooperative subjects, and high spectral quality. The variability of metabolite estimates that is introduced by the choice of analysis software is substantial, raising concerns about the robustness of MRS research findings, which typically use a single algorithm to draw inferences from much smaller sample sizes.

## Abbreviations

LCM: linear-combination modelling
tNAA: total N-acetylaspartate
tCho: total choline
mI: myo-Inositol
Glx: glutamate+glutamine
tCr: total creatine
MM: macromolecular
HSVD: Hankel singular value decomposition
CV: coefficient of variation

## Acknowledgement

This work is supported by NIH grants R01 EB016089 R01 EB023963 R21A G060245. GO receives support from NIH grant K99 AG062230. MP is supported by NIH grants P41EB015909 and R01NS106292.

The authors would like to thank Martin Wilson (University of Birmingham) for detailed explanations on Tarquin’s algorithm and Mark Mikkelsen (The Johns Hopkins University School of medicine) for setting up the linear mixed-effect model.

## Supplementary Material

**Supplementary Material 1.**
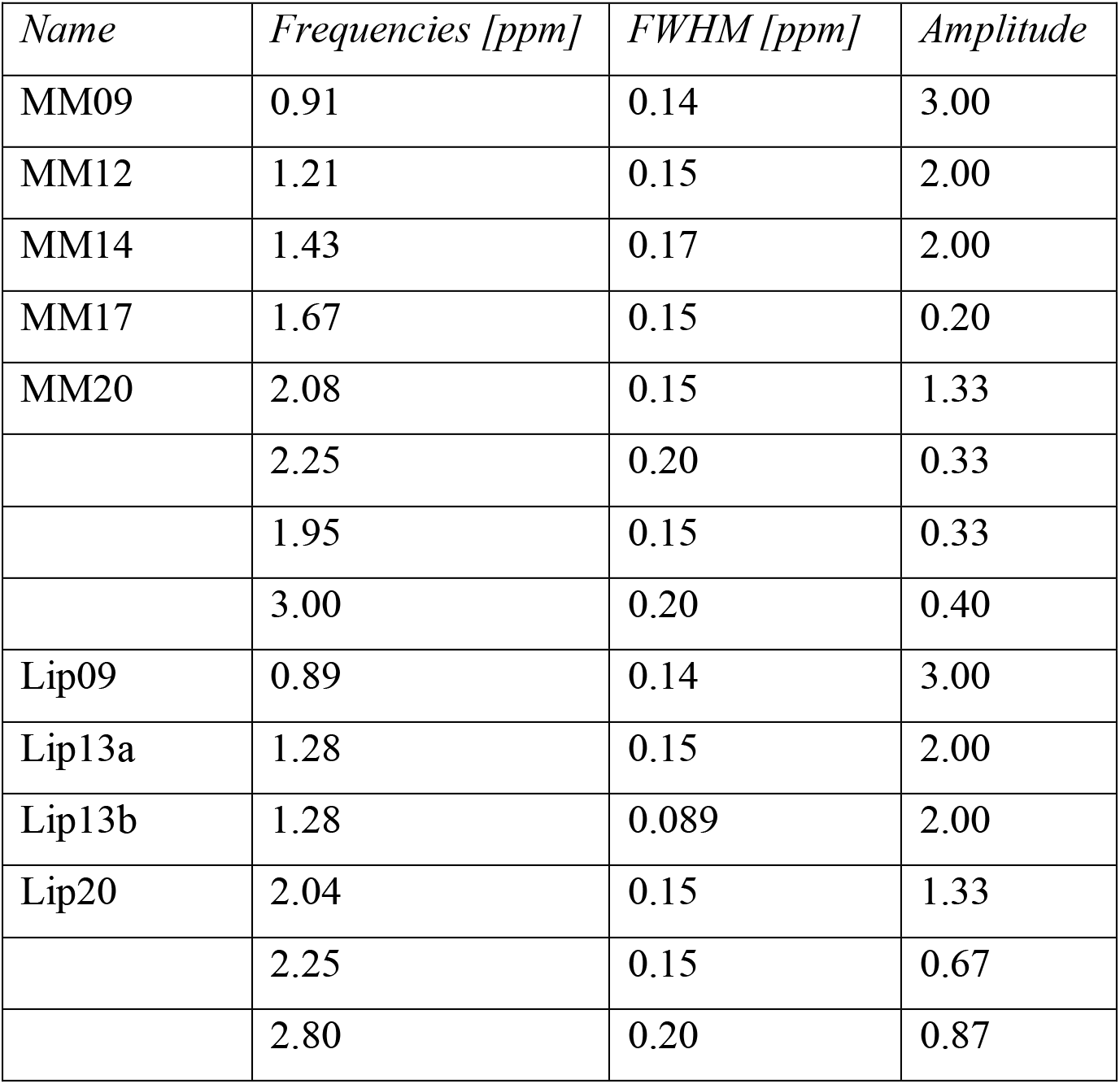
Properties of the Gaussian functions of the broad macromolecule and lipid resonances included in the basis sets, taken from section 11.7 of the LCModel manual. The amplitude values are scaled relative to the CH_3_ singlet of creatine with amplitude 3.

**Supplementary Material 2.**
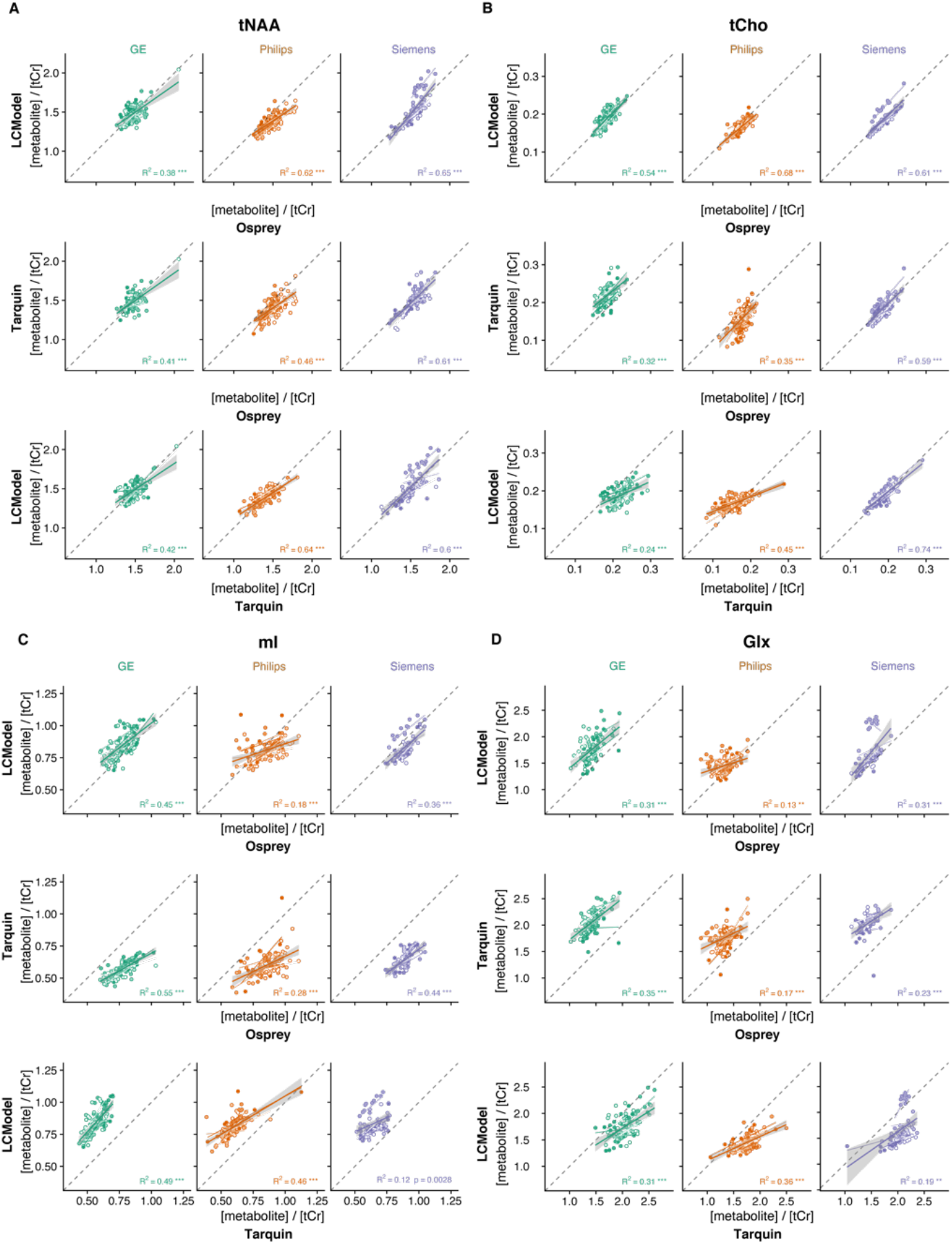
Facetted pair-wise correlational comparison of algorithms. LCModel and Osprey are compared in the first row, Tarquin and Osprey are compared in the second row, and LCModel and Tarquin are compared in the third row. Each sub-plot (A-D) corresponds to a different metabolite. Within-vendor (bold line with confidence interval) and within-site (thin line) correlations are color-coded. Asterisks indicate significant correlations (adjusted p < 0.01 = ** and adjusted p < 0.001 = ***).

**Supplementary Material 3.**
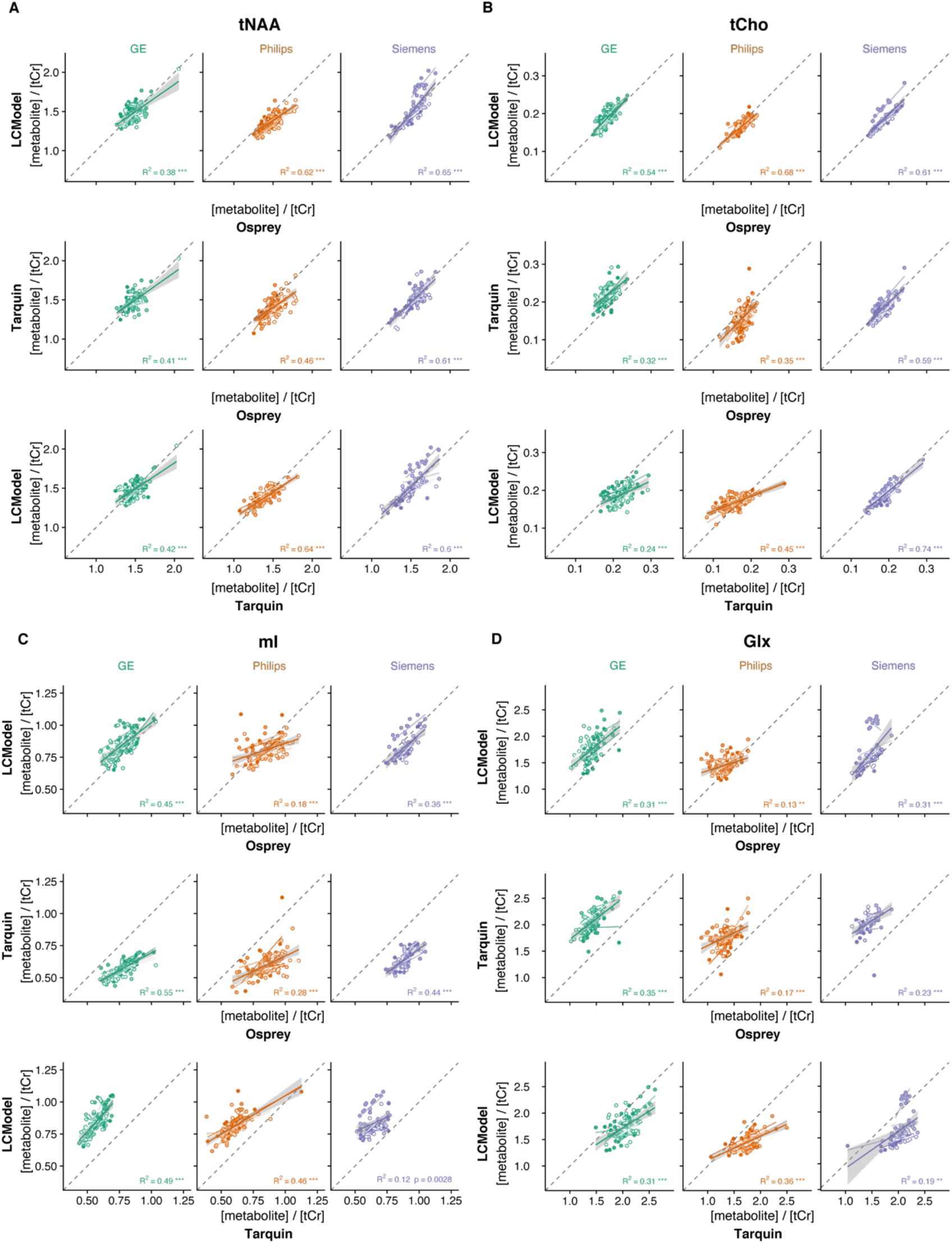
Variability of tCr models. Mean models +/- standard deviation (shaded areas) are presented column-wise by vendor and color-coded by LCM algorithm. The distribution and CV of the areas under the models are inset (A).Distribution and CV of the water referenced tCr concentrations are presented column-wise by vendor and color-coded by LCM algorithm.

